# Paradoxical hyperexcitability from Na_V_1.2 sodium channel loss in neocortical pyramidal cells

**DOI:** 10.1101/2021.02.02.429423

**Authors:** Perry W.E. Spratt, Roy Ben-Shalom, Atehsa Sahagun, Caroline M. Keeshen, Stephan J. Sanders, Kevin J. Bender

## Abstract

Loss-of-function variants in the gene *SCN2A*, which encodes the sodium channel Na_V_1.2, are strongly associated with autism spectrum disorder and intellectual disability. An estimated 20-30% of children with these variants are co-morbid for epilepsy, with altered neuronal activity originating in neocortex, a region where Na_V_1.2 channels are expressed predominantly in excitatory pyramidal cells. This is paradoxical, as sodium channel loss in excitatory cells would be expected to dampen neocortical activity rather than promote seizure. Here, we examined pyramidal neurons lacking Na_V_1.2 channels and found that they were intrinsically hyperexcitable, firing high-frequency bursts of action potentials (APs) despite decrements in AP size and speed. Compartmental modeling and dynamic clamp recordings revealed that Na_V_1.2 loss prevented potassium channels from properly repolarizing neurons between APs, increasing overall excitability by allowing neurons to reach threshold for subsequent APs more rapidly. This cell-intrinsic mechanism may therefore account for why *SCN2A* loss-of-function can paradoxically promote seizure.

## Introduction

Genetic variation in *SCN2A* is major risk factor for neurodevelopmental disorders, including various developmental epilepsies, autism spectrum disorder and intellectual disability. *SCN2A* encodes Na_V_1.2, a voltage-gated sodium channel that supports neuronal excitability throughout the brain, including cortical regions where it is expressed primarily in excitatory neurons (Hu et al., 2009; Spratt et al., 2019). Consistent with this expression pattern, *SCN2A* variants that enhance Na_V_1.2 function, and therefore excitatory activity in cortex, are most commonly associated with epilepsy. By contrast, loss-of-function (LoF) variants that dampen or eliminate channel function are typically associated with intellectual disability and autism spectrum disorder (ASD) (Howell et al., 2015; Sanders et al., 2018; Wolff et al., 2017). And yet, an estimated 20-30% of children with *SCN2A* LoF variants develop epilepsy (Sanders et al., 2018). The cellular mechanisms underlying seizure in conditions where sodium channel impairments are largely restricted to excitatory neurons is unknown.

Mouse models of heterozygous *Scn2a* loss (*Scn2a*^*+/-*^) suggest that seizures originate in neocortical pyramidal cells (Miyamoto et al., 2019; Ogiwara et al., 2018). Neocortical electroencephalograms (EEGs) from *Scn2a*^*+/-*^ mice exhibit spike-and-wave discharges characteristic of absence epilepsies at relatively low frequency (<10 events per hour) (Ogiwara et al., 2018). Similar EEG patterns have been observed in mice with conditional heterozygous expression of *Scn2a* in excitatory, but not inhibitory neocortical neurons (Ogiwara et al., 2018). However, changes in action potential (AP) output, a feature common to many models of Na_V_-channelopathy-mediated epilepsy (Goff and Goldberg, 2019; Li et al., 2020; Lopez-Santiago et al., 2017; Martin et al., 2010; Tai et al., 2014), have not been observed in mature *Scn2a*^*+/-*^ pyramidal cells (Shin et al., 2019; Spratt et al., 2019).

Given the incomplete penetrance of seizure in children with *SCN2A* LoF, as well as the low rates of spike-and-wave discharges observed in *Scn2a*^*+/-*^ mice, seizure-associated cellular phenotypes may be too subtle to discern in heterozygotes. In such cases, homozygous deletion may help identify cellular mechanisms that are disrupted by gene loss, as more overt effects may be revealed in a gene dose-dependent manner. This appears to be the case for *Scn2a* LoF conditions, as full deletion of *Scn2a* from striatally-projecting layer 5b pyramidal cells alone can recapitulate spike-and-wave discharge phenotypes observed in mice heterozygous for *Scn2a* in all pyramidal cells (Miyamoto et al., 2019). Thus, understanding how complete loss of *Scn2a* in layer 5b pyramidal cells results in hyperexcitability may shed light on seizure susceptibility in general.

Here, we examined cell-autonomous features of neuronal excitability in mice with conditional deletion of *Scn2a* in prefrontal pyramidal cells. Remarkably, despite deletion of a sodium conductance, neurons were more excitable, exhibiting both an increase in overall action potential output and an increase in high-frequency AP bursts. This was due to the unique function and localization of Na_V_1.2 in pyramidal cell neuronal compartments relative to other sodium channel and potassium channel classes. Na_V_1.2 was not responsible for action potential initiation, a process dependent instead on Na_V_1.6 channels in the axon initial segment (AIS) (Hu et al., 2009). Rather, Na_V_1.2 was critical for propagating action potentials through the soma and dendrites, and for activating voltage-gated potassium channels that repolarized neurons. Loss of Na_V_1.2 resulted in an increase in neuronal excitability largely through a failure to properly repolarize neurons between action potentials, and aspects of normal excitability could be rescued by injection of Na_V_1.2 conductance via dynamic clamp. Thus, the interplay between ion channel classes can affect electrogenesis in counterintuitive ways, highlighting the importance of considering neuronal excitability in different cellular compartments in neurodevelopmental channelopathy conditions.

## Results

### Hyperexcitability of AP initiation from Nav 1.2 deletion

Though *Scn2a*^*+/-*^mice exhibit spike-and-wave discharge epileptiform activity (Ogiwara et al., 2018), *ex vivo* studies of *Scn2a*^*+/-*^ pyramidal cells have yet to identify cellular mechanisms that could explain such events, including possible cell-intrinsic hyperexcitability. In fact, in the first postnatal week, pyramidal cells are less excitable, due to loss of Na_V_1.2 in axonal regions (Gazina et al., 2015; Spratt et al., 2019). Here, we hypothesized that complete homozygous loss of *Scn2a* may uncover dose-dependent excitability phenotypes related to Na_V_1.2 loss that are more difficult to observe in heterozygotes.

To examine neuronal excitability in the absence of Na_V_1.2 channels, we used a mouse line homozygous for conditional *Scn2a* knockout under Cre-recombinase control (*Scn2a*^*fl/fl*^). AAV-EF1α-Cre-mCherry was injected unilaterally into the medial prefrontal cortex (PFC) of *Scn2a*^*fl/fl*^ or *Scn2a*^*+/fl*^ mice at postnatal day 28-44 and neuronal excitability was examined in *ex vivo* acute slices following >4 weeks of viral expression. Na_v_ 1.2 deletion in *Scn2a*^*fl/fl*^-injected cells was confirmed with immunofluorescent staining of Na_V_1.2 in the AIS, comparing infected vs. uninfected hemispheres (**Fig. S1A-B**). Whole-cell current clamp recordings were made from mCherry positive L5b neurons in the injected hemisphere and compared to mCherry negative neurons in the contralateral uninfected hemisphere, or neurons in age-matched wild type (WT) mice. For simplicity, neurons conditionally lacking one or both *Scn2a* alleles are termed *Scn2a*^*+/-*^ and *Scn2a*^*-/-*^ throughout the manuscript, whereas mCherry negative neurons from either *Scn2a*^*+/fl*^ or *Scn2a*^*fl/fl*^ animals are pooled and termed WT.

Action potential excitability was assessed with somatic current steps of increasing intensity (Firing/Current [F/I] curves, **Fig. 1A-B**). Consistent with previous findings, the F/I curves of *Scn2a*^*+/-*^ neurons were no different than WT (*Scn2a*^*+/+*^: 3.78 ± 0.13 APs per 100 pA between 100–300 pA, n = 77; *Scn2a*^*+/-*^: 3.54 ± 0.21, n = 40, p = 0.193); however, *Scn2a*^*-/-*^ neurons exhibited a pronounced increase in F/I slope (**Fig. 1B**; *Scn2a*^*-/-*^: 4.55 ± 0.13, n = 60; *Scn2a*^*+/+*^ vs. *Scn2a*^*-/-*^ and *Scn2a*^*+/-*^ vs. *Scn2a*^*- /-*^: p < 0.001). This was due to both an increase in the number of APs generated during high-frequency bursts at current onset as well as an increase in steady state AP number closer to current offset in *Scn2a*^*--l*^ neurons (**Fig. 1B**).

**Figure 1:**
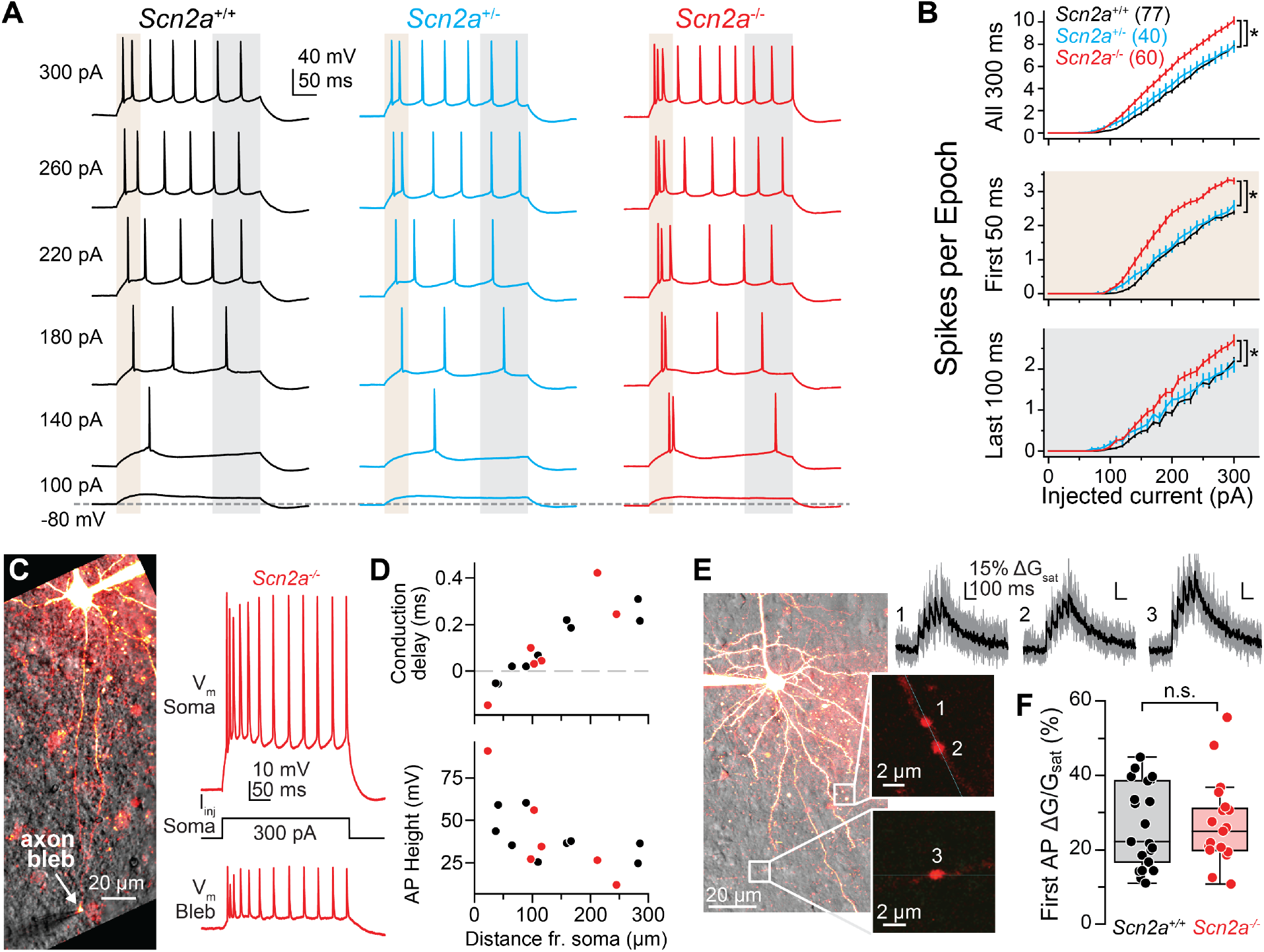
*Scn2a* knockout increases AP excitability, with intact axonal propagation. **A:** APs (spikes) per 300 ms stimulation epoch for each current amplitude in *Scn2a*^*+/+*^ (black), *Scn2a*^*+/-*^ (cyan), and *Scn2a*^*-/-*^ (red) cells. Brown and grey areas highlight first 50 ms and last 100 ms of epoch, respectively. Note increased burst firing and overall firing rates in *Scn2a*^*-/-*^ conditions. **B:** AP number vs current injection curves for 3 *Scn2a* conditions, color coded as in A. Bars are mean ± SEM. n denoted in parenthesis. Top: entire 300 ms epoch. Middle: first 50 ms. Bottom: last 100 ms. *: FI slope between 100 and 300pA, p< 0.01, Kruskal-Wallis test **C:** Scanning DIC and 2-photon microscopy fluorescent image of simultaneous recording in a *Scn2a*^-/-^ pyramidal neuron of APs evoked from soma and recorded at axon bleb 212 µm from axon hillock. **D:** Conduction delay of AP threshold between soma and bleb in each recording vs. axon length and AP amplitude in bleb in *Scn2a*^*+/+*^ (black) and *Scn2a*^*-/-*^ (red) cells. **E:** Scanning DIC and 2-photon microscopy fluorescent image detailing bouton scan sites for AP-evoked calcium imaging of a *Scn2a*^-/-^ neuron (insets). Responses to burst of 5 APs at boutons 1-3 shown as average (black) overlaid on single trials (grey, 10 trials total). First AP amplitude is no different between *Scn2a*^*+/+*^ and *Scn2a*^*-/-*^ cells.

To better understand how neuronal excitability is affected by *Scn2a* loss, we first examined the expression and function of Na_V_1.6, which is expressed in addition to Na_V_1.2 in mature neocortical pyramidal cells. In contrast to Na_V_1.2, which is localized primarily to somatodendritic compartments, Na_V_1.6 is expressed at high levels in the axon and mediates the initiation and propagation of orthodromic APs (Hallermann et al., 2012; Hu et al., 2009; Li et al., 2014; Spratt et al., 2019). To assay Na_V_1.6 function, we first measured persistent Na_V_ currents generated from voltage steps from -90 to -60 or -55 mV, as these currents reflect the recruitment of low-threshold AIS-localized Na_V_s (Taddese and Bean, 2002). Furthermore, we examined the intensity of Na_V_1.6 immunofluorescent staining in the AIS in infected versus uninfected hemispheres. In both cases, modest increases in channel function/expression were evident: median persistent currents were increased by 18% for steps from -90 to -60 mV in *Scn2a*^-/-^ neurons (**Fig. S1G-H**), and Nav 1.6 fluorescent intensity, normalized to its AIS scaffolding partner, ankyrin-G, was increased by 13% (**Fig S1A-D**).

We examined other aspects of neuronal excitability, including the resting membrane potential, input resistance and whole-cell potassium currents. Resting membrane potential did not differ with *Scn2a* expression levels (*Scn2a*^*+/+*^: 74.9 ± 0.4 mV, n = 92; *Scn2a*^*+/-*^: 73.8 ± 1.3, n = 47, *Scn2a*^*/-*^: 73.9 ± 0.6, n = 47, p = 0.177). As with previous comparisons between WT and *Scn2a*^*+/-*^ cells (Spratt et al., 2019), potassium currents were not different between WT and *Scn2a*^*-/-*^ cells (**Fig. S1I-J**). Input resistance, however, was higher in *Scn2a*^*-/-*^ cells, potentially contributing to increased excitability. To test this, we compared F/I slope to neuronal input resistance (**Fig. S1K-L**). While a modest correlation was observed in WT cells (R^2^ = 0.14), no such correlation was observed in *Scn2a*^*+/-*^ or *Scn2a*^*-/-*^ cells (R^2^ = 0.006 and 0.004, respectively. p > 0.5), where slopes were higher even in cells with low input resistance. This suggests that increased AP output is unlikely to be explained by modest changes in intrinsic membrane resistance alone.

In light of these observations, we predicted that axonal AP propagation, supported by Na_V_1.6, was likely to be intact, even with Na_V_1.2 loss. To test this, we first made simultaneous whole-cell recordings from the soma and an axonal bleb, formed during slice preparation as the primary axon is transected at the slice surface. Blebs were identified at various distances from the axon hillock in both WT and Cre+ *Scn2a*^*-/-*^ cells, and APs were evoked via somatic current injection. Conduction delays between soma and bleb were unaltered when accounting for axon length (**Fig. 1C-D**; Conduction velocity: WT: 1.26 ± 0.29 m/s, n = 7; -/-: 1.21 ± 0.36, n = 5, p = 0.8 Mann-Whitney). Next, we confirmed that APs could propagate to neurotransmitter release sites by measuring AP-evoked calcium transients in axonal boutons using 2-photon microscopy. Here, APs reliably generated calcium transients in *Scn2a*^*-/-*^ boutons, with no differences in transient amplitude compared to WT boutons (**Fig. 1E**). Together, these data indicate that axonal AP propagation is intact in *Scn2a*^*-/-*^ cells, likely due to continued expression of axonal Na_V_1.6 channels.

### Hypoexcitability of dendritic AP backpropagation from Na_V_1.2 deletion

Heterozygous loss of Na_V_1.2 impairs somatodendritic excitability in mature neocortical pyramidal cells, affecting both action potential waveform recordings at the soma and AP-evoked calcium signaling in dendrites (Spratt et al., 2019). To determine the effects of full Na_V_1.2 deletion, we examined AP waveform at the soma using phase-plane analysis, which compares the rate of change in voltage during APs to the voltage (Jenerick, 1963). These plots help reveal different aspects of AP initiation, including a sharp kink at AP threshold and two distinct components of the rising phase of an AP that reflect the initial activation of AIS Na_V_s and subsequent recruitment of somatodendritic Na_V_s (**Fig. 2B-C**). Analysis of APs evoked at rheobase showed that AP threshold was not altered by heterozygous or homozygous loss of *Scn2a* (**Fig. 2A-D**). Similarly, the AIS-associated component of the AP rising phase was unaltered (**Fig. 2C-D**). Backpropagation of this AIS-evoked AP into the soma appeared to be reliable, as assayed by evoking APs from a whole-cell recording of a near-AIS axonal bleb while monitoring somatic voltage (**Fig. S2**). Together, these data are consistent with normal AIS AP initiation, mediated by Na_V_1.6 channels.

**Figure 2:**
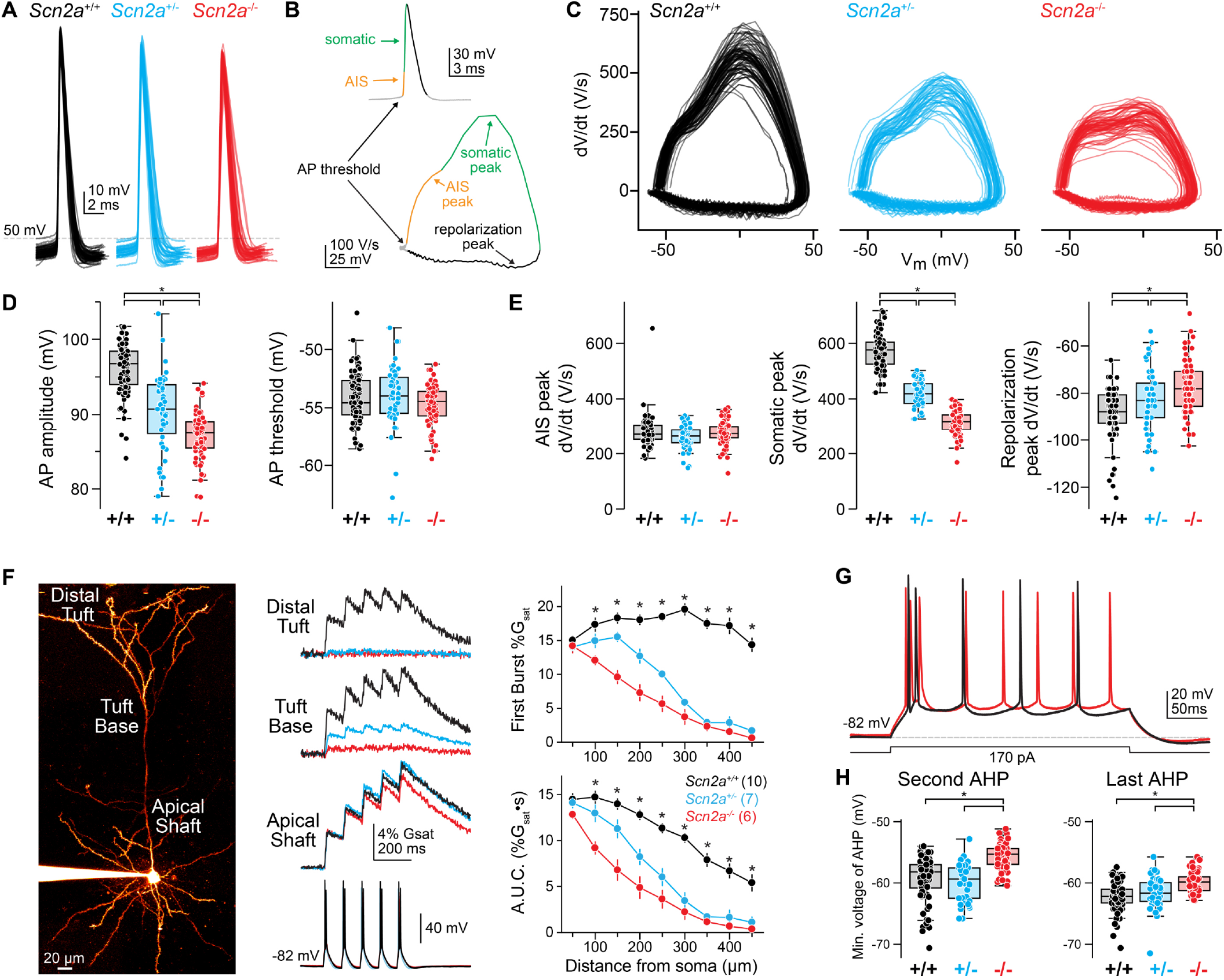
*Scn2a* knockout impairs somatodendritic excitability. **A:** Rheobase AP waveform from all neurons studied. Note progressive reduction in AP height with loss of Na_V_1.2 expression. **B:** AP plotted as voltage vs. time (top) and dV/dt vs. voltage (phase-plane plot, bottom). Different phases of the AP are color-coded across panels to indicated different phases of the AP corresponding to initiation of AP in AIS, the soma, and peak repolarization. **C:** Phase-plane plots of data shown in (A). **D:** AP amplitude and threshold for APs at rheobase. Circles are single cells. Box plots are median, quartiles, and 90% tails. Data color coded as in (A). *: p < 0.001, Kruskal-Wallis test. **E:** Peak of AIS and somatic components of the rising phase of the AP, and minimum of the falling phase of the AP. *: p < 0.001, Kruskal-Wallis test. **F:** Left, Morphology of imaged neuron. Middle, 2PLSM calcium imaging of AP-evoked calcium transients throughout apical tuft dendrites in *Scn2a*^*+/+*^ (black), *Scn2a*^*+/-*^ (cyan), and *Scn2a*^*-/-*^ (red) cells, n denoted in parenthesis. Right, transient amplitude shown for first of 5 bursts (top) and area under the curve from stimulus onset to stimulus offset +100 ms (bottom). Circles and bars are means ± SEM. *: p < 0.01, Kruskal-Wallis test. **G:** Overlaid AP response to 170 pA current injection in *Scn2a*^*+/+*^ (black) and *Scn2a*^*-/-*^ (red) cell. Note difference in AHP amplitude. **H:** AHP amplitude of the second (left) and final (right) interspike intervals during spike trains elicited by 300pA current injection. *: p < 0.001, Kruskal-Wallis test.

While axonal aspects of AP initiation were not affected by *Scn2a* loss, somatodendritic features of AP initiation were increasingly impaired as *Scn2a* expression was reduced to 50% (heterozygote) or eliminated entirely (conditional knockout). AP height and the speed of depolarization during the somatodendritic component of the AP rising phase both reflect engagement of somatodendritic sodium channels by APs as they backpropagate from the AIS to the soma. Consistent with previous observations (Spratt et al., 2019), these features were smaller than WT in *Scn2a*^*+/-*^ cells and smaller still in *Scn2a*^*-/-*^ cells (**Fig. 2A-E**). Similarly, AP-evoked dendritic calcium transients, which reflect local dendritic activation of voltage-gated calcium channels, were impaired markedly in *Scn2a*^*-/-*^ cells (**Fig. 2F**). Thus, axonal and dendritic function are differentially affected by Na_V_1.2 deletion, with increased excitability in measures of AP output, and decreased excitability in measures of dendritic function.

### Na_V_1.2 deletion affects potassium channel-mediated AP repolarization

Just as membrane potential fluctuations preceding an AP affect AP threshold (Azouz and Gray, 2000; Bender and Trussell, 2009; Hu et al., 2009; Kole et al., 2007; Uebachs et al., 2006), the amplitude of the AP affects its repolarization by changing the driving force on potassium channels. Indeed, in *Scn2a*^*-/-*^ cells, where AP height is reduced by 9 mV, we found that AP repolarization speed was reduced 12% (WT: -88.69 ± 1.14 mV, n = 91 vs *Scn2a*^*-/-*^: -77.38 ± 1.35, n = 67) and that membrane potential between APs (e.g., afterhyperpolarization, AHP) was more depolarized than in WT cells throughout spike trains (2^nd^ AHP with 300pA current injection: WT: mean = -59.4, SEM = 0.41, n = 77 vs -/-: mean = -55.68, SEM = 0.28, n = 56, p < 0.001; final AHP: WT: mean = -62.28, SEM = 0.26, n = 77, -/-: mean = -59.96:, SEM = 0.24, n = 56, p<0.001). This effect may contribute to hyperexcitability phenotypes observed in cells with Na_v_ 1.2 deletion by allowing neurons to reach threshold for ^V^ ^V^ subsequent APs more quickly

To better understand how each of these effects affect neuronal excitability, we examined how Na_V_1.2 loss and modest increases in Na_V_1.6 affected repetitive AP activity in compartmental neuron models with Na_V_1.6 and Na_V_1.2 distributed as previous described (Ben-Shalom et al., 2017; Spratt et al., 2019). We started by isolating increases in Na_V_1.6 without altering Na_V_1.2 to determine the effect this single manipulation has on AP waveform and repetitive activity. Na_V_1.6 density was increased by 20% and 50%, matching or exceeding increased Na_V_1.6 staining and persistent sodium current observed in *Scn2a*^*-/-*^ cells (**Fig. S3**). As expected from increasing Na_V_ density, the speed of AP depolarization was increased; however, increasing NaV1.6 density reduced, rather than increased, the frequency of subsequent APs. Thus, modest increases in Na_V_1.6 appears unable to account for hyperexcitability observed in *Scn2a*^*-/-*^ conditions. Instead, models were able to capture several aspects present in empirical data simply by lowering Na_V_1.2 density. Excitability increased only modestly at 50% Na_V_1.2 density, but increased dramatically with complete Na_V_1.2 removal (**Fig. 3A**), mirroring effects observed in *Scn2a* and *Scn2a*^*-/-*^ conditions (**Fig. 1**). Effects on AP output were similar whether or not Na_V_1.6 density was increased, suggesting that changes in Na_V_1.6 density have little effect on repetitive activity.

**Figure 3:**
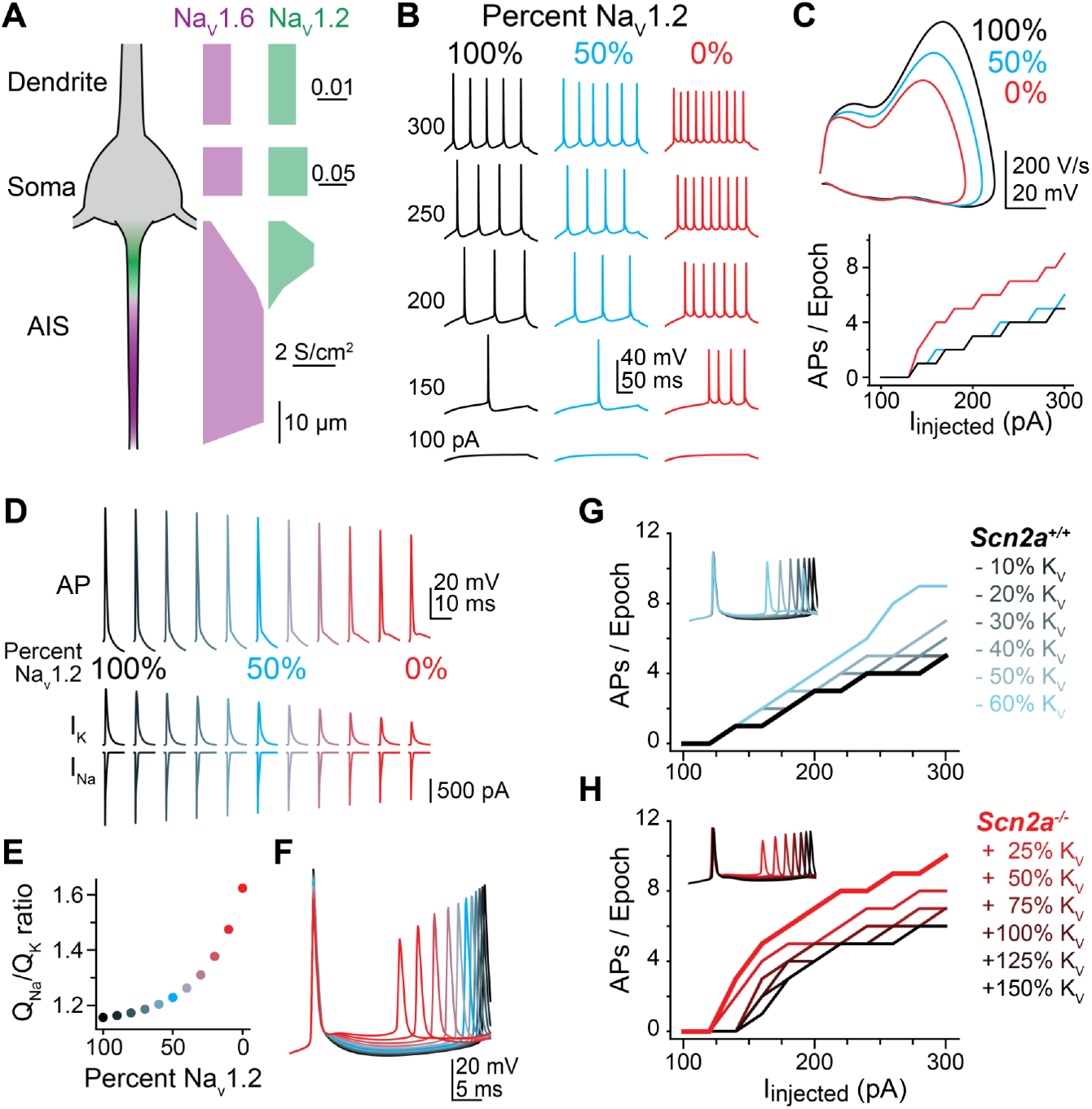
*Scn2a* loss in dendrites increases AP excitability in compartmental models. **A:** Compartmental model of cortical layer 5 pyramidal cell with Na_V_1.2 and Na_V_1.6 distributed in the AIS soma and dendrites as shown. Purple and green for Na_V_1.2 and Na_V_1.6 show relative densities in dendrite, soma, and AIS in model. Note different scale bars for each compartment. **B:** Example firing patterns of models in which Na_V_1.2 is reduced to 50 or 0% in all compartments. Data color coded as in Figs. 1-2. **C:** Phase-plane plots and F/I curves for each model. Note increase in F/I is appreciable only in *Scn2a*^*-/-*^ conditions. No other channels or model parameters altered except for Na_V_1.2 density. **D:** Plots of APs elicited by 350 pA current injection and underlying whole-cell sodium and potassium currents with progressive loss of Na_V_1.2 **E:** Charge transfer (Q) ratio (Na/K) for each AP as a function of Na_V_1.2 density. 100% is WT levels, 0 is *Scn2a*^*-/-*^. **F:** Timing of first APs in conditions noted in D-E (identical color-coding). Note depolarization of AHP and advanced AP2 timing with Na_V_1.2 loss. **G:** Reducing potassium conductance in WT model recapitulates hyperexcitability observed in *Scn2a*^*-/-*^ cells due to depolarization of AHP (inset). **H:** Increasing potassium conductance in *Scn2a*^*-/-*^ model reduces excitability, in part by delaying AP timing (inset).

As in neurons, effects on repetitive AP activity correlated with changes in AP height and potassium channel engagement. We therefore asked how potassium channels that are activated during AP repolarization were affected by progressive removal of Na_V_1.2 from our models. Interestingly, the relationship between Na_V_ and potassium channel current generated during an AP, as Na_V_1.2 density was reduced, was exponential rather than linear. At 50% Na_V_1.2 density, the ratio of the charge transfer (Q) of sodium vs. potassium current during the AP was only 6% higher than in 100% Na_V_1.2 density conditions. By contrast, this ratio was 40% higher when Na_V_1.2 was deleted completely (**Fig. 3D-F**). This exponential relationship may account for why *Scn2a* heterozygotes have F/I curves that are comparable to WTs, whereas marked hyperexcitability is clear in *Scn2a*^*-/-*^ cells. *Scn2a* cells.

This modeling suggests that the interplay between Na_V_1.2 and voltage-gated potassium channels, rather than changes in Na_V_1.6 density, are the primary drivers of hyperexcitability in *Scn2a*^*-/-*^ cells Consistent with this, increased potassium channel density in *Scn2a*^-/-^ models reduces excitability, while reduced potassium channel density in *Scn2a*^+/+^ models increases excitability (**Fig. 3G-H**). Together, these models predict that injection of conductances that mimic Na_V_1.2 engaged during AP depolarization, or potassium channels engaged during AP repolarization, should restore features of WT excitability in *Scn2a*^*-/-*^ cells. We tested this using somatic dynamic clamp injection of Nav 1.2-like or Kv 1.2-like conductances into *Scn2a*^*-/-*^ cells interleaving epochs with and without dynamic clamp in all conditions (**Fig. 4A-B**). Kv1.2 was chosen since it is commonly expressed in neocortical pyramidal cells and helps control repetitive firing frequency (Guan et al., 2007a). For Na_V_1.2 conductance injection, injection amplitude was adjusted to restore peak dV/dt of the AP to 88% of WT levels in these recording conditions (see Methods) without inducing feedback within the amplifier circuitry (WT: 732 ± 26 V/s, n = 5; -/-baseline: 412 ± 12, post 0.75–1.5 µS injection: 646 ± 24, n = 9). K_V_1.2 conductance injection, as expected, had no effect on peak dV/dt. Therefore, we examined two different injection intensities in each cell (100–200 nS), as these appeared to encompass a range of excitability effects similar to Na_V_1.2 conductance injection.

**Figure 4:**
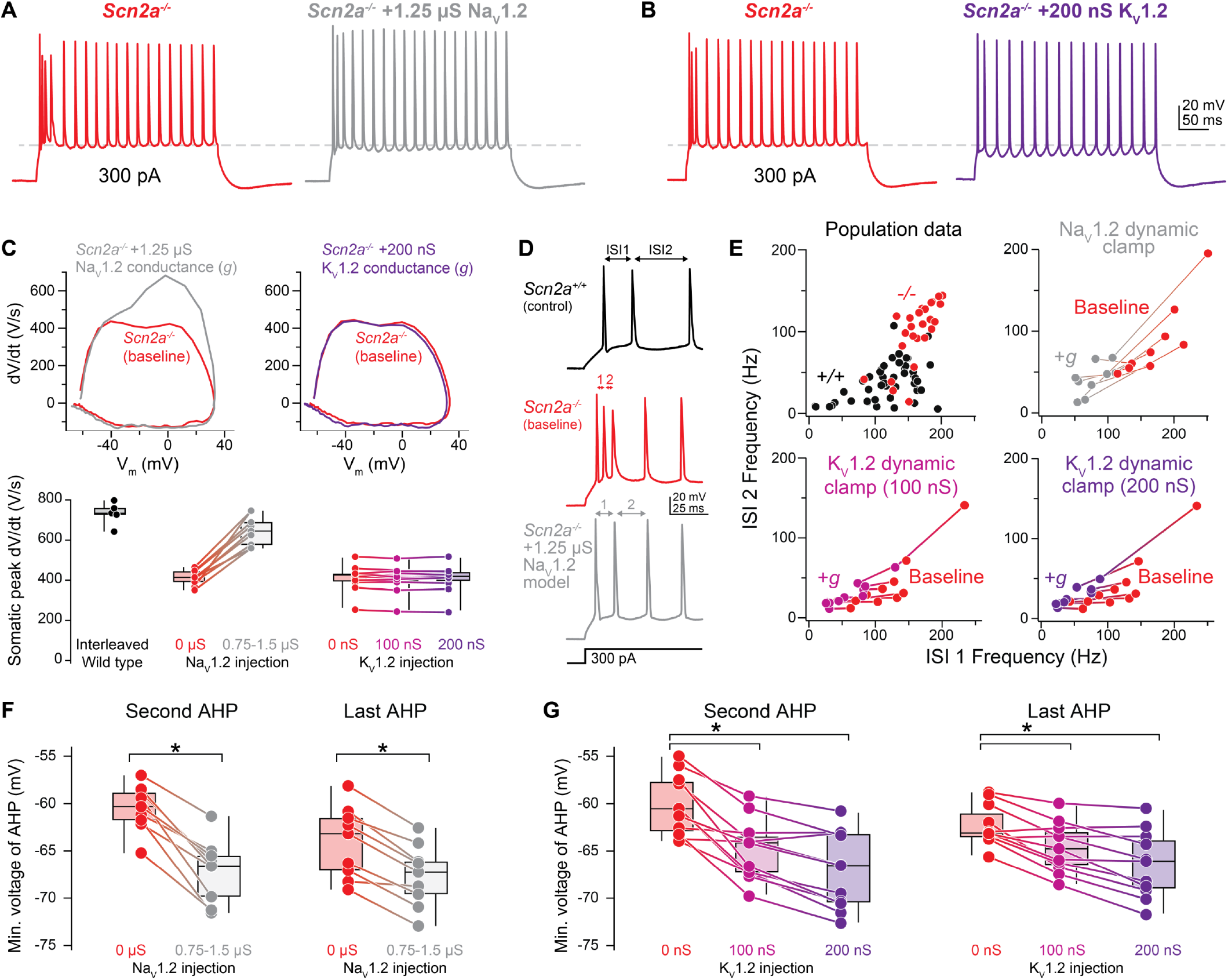
Dynamic clamp injection of Na_V_1.2 or K_V_1.2 conductance rescues excitability. **A:** AP firing response to 300 pA, 300 ms current injection before (red) and after (grey) injection of 1.25 µS Na_V_1.2 conductance via somatic recording electrode. Dashed line aligned to last AHP in baseline conditions. **B:** Same as A, but for injection of 200 nS of K_V_1.2. Baseline in red, K_V_1.2 injection in purple. **C:** Top, phase planes of first AP for each condition in A-B. Bottom, summary data for interleaved WT cells in identical recording conditions and for all *Scn2a*^*-/-*^ cells with Na 1.2 or K 1.2 conductance injection. Circles are single cells with lines connecting paired data. **D:** Highlight of timing of first 3 APs in WT and *Scn2a*^*-/-*^ before and after Na_V_1.2 conductance injection. Inter-spike intervals (ISIs) for APs 1-2 and 2-3 plotted in E. **E:** Top left, ISI1 vs. ISI2 at 300 pA current injection for all WT (black) and *Scn2a*^*-/-*^ (red) cells in population in Fig. 1. Other panels show change in ISIs from *Scn2a*^*-/-*^ baseline (red) after conductance injection (other colors). Lines connect values from the same cell. **F:** Minimum voltage between APs (e.g., AHP) between APs 1-2 and the penultimate and last APs for 300 pA stimulus injection before and after Na_V_1.2 conductance injection. Data shown as in C. *: p < 0.01, Wilcoxon signed rank test. **G:** Same as F, but for injection of 100 and 200 nS of K_V_1.2.

*Scn2a*^*-/-*^ cells often fire high frequency bursts at current onset and have AHPs that are more depolarized than observed in WT cells (**Fig. 1A-B, 2G**). Injection of either Nav 1.2- or. K_V_1.2 like conductances suppressed burst generation, as assayed by the changes in instantaneous frequencies of APs 1-2 and 2-3 within spike trains (**Fig. 4D-E**). Similarly, AHP amplitudes were hyperpolarized by injection of either Na_V_1.2 or K_V_1.2 (**Fig. 4F-G**). But despite restoring these aspects of neuronal excitability to WT-like levels, and despite observing changes in the first 50 ms of F/I curves, overall F/I curves (300 ms current injection epochs) were largely unaffected by Na_V_1.2 or Kv1.2 conductance injection (**Fig. S4A**). This may be due to the spatial extent over which conductances, injected via a somatic electrode, can influence the neuronal membrane potential. Transient conductances, including Na_V_1.2 or K_V_1.2-like conductances injected here, may have a large effect on somatic membrane potential but limited effect on the axonal and dendritic membrane potential, due to difficulties in rapidly charging membranes more distal to the injection electrode, akin to space-clamp errors in voltage-clamp recordings (Williams and Mitchell, 2008). We tested this idea *in silico* by examining F/I curves in cases where Na_V_1.2 was rescued only in the soma and/or proximal AIS compared to cases where Nav1.2 was restored in dendritic compartments as well (**Fig. S4B**). As in our dynamic clamp experiments, we found that restoration just in the soma or AIS had little effect on overall F/I curves, even when Nav1.2 density was restored to 80-90% of WT levels (e.g., similar to dynamic clamp rescue of dV/dt). This suggests that dendritic interactions between Nav1.2 and potassium channels are critical in regulating overall pyramidal cell excitability and highlights the importance of Na_V_1.2 in dendritic excitability.

## Discussion

Neurodevelopmental epilepsies are associated with variation in all three central nervous system-localized sodium channels. For *SCN1A*, epilepsy stems almost exclusively from variants that impair channel function (e.g., loss-of-function), due to resulting deficits in inhibitory interneuron excitability and circuit disinhibition (Goff and Goldberg, 2019; Tai et al., 2014). For *SCN8A*, epilepsy manifests largely from gain-of-function variants that produce neuronal hyperexcitability in excitatory networks (Lopez-Santiago et al., 2017). *SCN2A* is unique in that both gain- and LoF are associated with seizure. For gain-of-function *SCN2A* variants, where the age of seizure onset is within days to months of birth, the underlying mechanisms appear related to hyperexcitability in glutamatergic neuronal axons (Gazina et al., 2015; Sanders et al., 2018). For LoF conditions, where the age of seizure onset is typically after 12 months of life (Brunklaus et al., 2020), the underlying cellular mechanisms have been less clear. Here, we identified a candidate mechanism for such hyperexcitability: the interplay between sodium and potassium channel electrogenesis and the resulting increased excitability due to a failure to properly repolarize neurons between APs.

Several lines of evidence support this hypothesis. First, *in vivo* data from rodent models indicates that aberrant EEG signatures in *Scn2a* LoF conditions arise from dysfunction in neocortical pyramidal cells (Miyamoto et al., 2019; Ogiwara et al., 2018) and that conditional heterozygous expression of *Scn2a* in neocortical inhibitory neurons has no effect on EEG patterns (Ogiwara et al., 2018). Furthermore, complete deletion of *Scn2a* exclusively from striatally projecting layer 5b pyramidal cells has similar effects on EEG activity. Together, these data indicate that dysfunction in layer 5 pyramidal cell physiology is likely causal to seizure phenotypes.

Second, the timing of seizure onset in LoF conditions corresponds with a developmental switch from Na_V_1.2 to Na_V_1.6 in the distal AIS (Gazina et al., 2015; Hu et al., 2009; Spratt et al., 2019). Before this switch, *SCN2A* LoF would impair AP electrogenesis. But after this switch, Na_V_1.2 has no role in AP initiation. Indeed, both AP threshold and the AIS component of the AP waveform are unaffected in conditional *Scn2a* heterozygous or homozygoous knockout neurons (**Fig. 2**). Instead, Na_V_1.2 becomes critical for dendritic electrogenesis, and we show here that progressive loss of Na_V_1.2 in neocortical dendrites affects AP repolarization and, therefore, can advance the timing of subsequent APs. This separation of roles for different Na_V_ genes—with Na_V_1.6 governing axonal electrogenesis and Na_V_1.2 regulating dendritic electrogenesis— establishes, to our knowledge, the only condition in which LoF in a sodium channel could result in AP hyperexcitability. By contrast, LoF in *SCN1A* or *SCN8A* both result in hypoexcitability, as these channels remain localized to the AIS of neurons in which they are expressed (Katz et al., 2018; Makinson et al., 2017; Ogiwara et al., 2007).

Given these observations, why then is seizure associated with only an estimated 20-30% of those with *SCN2A* LoF variants? Moreover, if seizure is occurring on a heterozygous background in children, and can be detected at low frequency in EEG from heterozygous *Scn2a* mice, why is neuronal hyperexcitability observable only in neurons with complete *Scn2a* deletion? A possible explanation comes from our neuronal simulations, where we noted an exponential, rather than linear, relationship in the charge transfer ratio of sodium and potassium with progressive loss of Na_V_1.2 channels. This ratio increases only slightly from 100% to 50% Na_V_1.2 conditions, but then increases dramatically after 50% loss. Consistent with this, a gene-trap approach that reduces *Scn2a* expression by ∼75% also results in neuronal hyperexcitability (Zhang et al., co-submitted).

In this way, heterozygous *SCN2A*-loss appears to lie at an inflection point of the curve where other aspects of neuronal excitability, perhaps encoded by common genetic variation in the population or due to variation of gene expression levels during development, could push neuronal excitability toward either side of this inflection point, thereby promoting or protecting from seizure. It is well-established that common genetic variants play a major role in a range of neurodevelopmental disorders, including epilepsy (Leu et al., 2019; Speed et al., 2014) and autism spectrum disorder (Gaugler et al., 2014). Furthermore, genetic background is a well-defined determinant of seizure severity in several murine epilepsy (Gu et al., 2020; Kang et al., 2019), including *Scn2a*^*+/-*^ mice in which aberrant EEG signals can be observed in *Scn2a*^*+/-*^ mice on a pure C57Bl/6J background but not when *Scn2a*^*+/-*^ C57Bl/6J mice are crossed to Black Swiss mice (Mishra et al., 2017). Based on this, one prediction would be that seizure frequency or severity should be higher in children with protein truncating variants than those that exhibit partial LoF missense variants that temper, rather than completely eliminate, channel function. Testing this hypothesis will require deep genotype-phenotype analysis of a large number of *Scn2a* variants in parallel with biophysical characterization of channel variants and predicted effects on neuronal excitability in neuronal models (Ben-Shalom et al., 2017; Wolff et al., 2017), controlling for symptom-based ascertainment. Non-genetic factors, such as medications or supplements, may also play a role, and seizure onset in children with SCN2A-related neurodevelopmental disorders should prompt a careful examination of recent therapeutic changes.

### The role of Na_V_1.2 in pyramidal cell excitability

Based on immunostaining, Na_V_1.2 channels were previously shown to be expressed in a region of the AIS proximal to the soma of neocortical pyramidal cells (Hu et al., 2009). Here, they have been proposed to provide an electrical boost to APs as they backpropagate from the Na_V_1.6-rich distal AIS initiation site into the somatodendritic compartment. One prediction of this model is that APs would fail to backpropagate into the soma entirely in the absence of Na_V_1.2 in the proximal AIS. Instead, AIS-initiated APs would only produce “spikelets” in the soma, which reflect passive invasion of the AP into the soma (Hu et al., 2009). We tested this here and, largely, did not observe differences in axonal backpropagation efficacy from axon to soma between WT and *Scn2a*^*-/-*^ cells (**Fig. S3**). However, this failure to observe spikelets does not rule out this hypothesis, as APs may have been boosted instead by increased Na_V_1.6 density in the AIS of *Scn2a*^*-/-*^ cells. Moreover, spikelets may be generated more commonly in species with larger pyramidal neurons, including rat as observed originally (Hu et al., 2009). Mouse prefrontal pyramidal cells, by contrast, are smaller than in other species and smaller than in more caudal brain regions. Ideally, future development of a selective, voltage-independent pharmacological inhibitor of Na_V_1.2, independent of Na_V_1.6, will be necessary to test these ideas further.

Beyond the AIS, Na_V_1.2 appears to play a major role in somatodendritic excitability. As in previous work (Spratt et al., 2019), we found that compartmental models of pyramidal cell excitability were best fit by placing equal numbers of Na_V_1.2 and Na_V_1.6 in somatodendritic regions. Under these conditions, progressive loss of Na_V_1.2 increasingly impaired AP-evoked calcium transients throughout the apical dendritic arbor (**Fig. 2**).\ Interestingly, similar dendritic excitability deficits have not been observed with conditional Na_V_1.6 knockout, although compensation via Na_V_1.2 in the AIS of such neurons is evident (Katz et al., 2018). Thus, it remains possible that Na_V_1.6 has similar contributions to dendritic excitability as Na_V_1.2, but these contributions may be difficult to identify with genetic approaches due to compensation from other Na_V_ isoforms.

In dendrites, loss of Na_V_1.2 has dual effects, both decreasing dendritic excitability and, paradoxically, increasing overall AP output. While the former is a direct consequence of the loss of Na_V_-mediated excitability in dendritic compartments, we show here that the latter is more indirect. By decreasing overall depolarization during an AP in dendritic regions, loss of Na_V_1.2 reduces the activation of voltage-gated potassium channels, in turn increasing the amplitude of inter AP AHPs. This increases overall firing rate, as neurons can then reach threshold for subsequent APs more quickly. This contrasts with other mechanisms by which potassium channels support high-frequency firing. For example, K_V_3 channels are critical for generating extremely high-frequency firing in auditory brainstem, cerebellum and in fast-spiking interneurons. In these cells, K_V_ 3 channels promote rapid repolarization, allowing Na_V_s to recover from inactivation during repetitive activity (Erisir et al., 1999; Kaczmarek and Zhang, 2017; Lau et al., 2000; Macica et al., 2003; Zagha et al., 2008). These types of interactions between K_V_s and Na_V_s may account for aspects of hyperexcitability associated with gain-of-function variants in potassium channels (Niday and Tzingounis, 2018).

### Considerations for understanding and treating genetic epilepsies

Considerable effort has been made towards understanding the cells and circuits that are etiological to seizure in multiple genetic epilepsies (Feng et al., 2019). Epilepsy associated with *SCN2A* LoF highlights the need for not only considering cell types and circuits important for seizure, but also the subcellular distribution of associated proteins and their interactions within that subcellular ion channel network (Poolos and Johnston, 2012). Work here highlights how further blockade of Na_V_s with sodium channel-blocking anti-epileptics—already counter-indicated in *SCN2A* LoF cases—may increase seizure burden. Furthermore, this work identifies potential candidates for future therapeutic development. For example, development of activators of potassium channels preferentially localized to the somatodendritic domain may help counteract the loss of Na_V_1.2 (Guan et al., 2007b). Due to the exponential relationship between reduced Na_V_1.2 function and neuronal hyperexcitability, even modest changes in potassium channel function could have profound benefits for seizure control. Whether similar interactions occur with other epilepsy-associated genes (e.g., *KCNQ2/3, SCN1A*, etc.) remains largely unexplored. Such studies could help identify novel approaches to treating such disorders.

## Acknowledgments

We are grateful to members of the Bender lab for critically assessing this work and to members of the FamilieSCN2A Foundation for generous discussions. This research was supported by SFARI grants 513133 and 629287 (KJB), the Natural Sciences and Engineering Research Council (NSERC) of Canada PGS-D Scholarship (PWES), and NIH grants NS095580 (RBS) and MH111662 (SJS).

## Author contributions

Conceptualization: PWES, RBS, KJB; Methodology: PWES, RBS, AS, CMK, KJB; Software: PWES, RBS; Formal Analysis: PWES, KJB; Investigation: PWES, RBS, AS, CMK, KJB; Resources: SJS, KJB; Writing—original draft: PWES, KJB; Writing—review & editing: all authors; Visualization: PWES, KJB; Supervision: KJB; Project Administration: KJB; Funding Acquisition: PWES, RBS, SJS, KJB.

## Materials and Methods

All experimental procedures were performed in accordance with UCSF IACUC guidelines. All experiments were performed on mice housed under standard conditions with ad libitum access to food and water. C57B6J (JAX: 000664) were obtained from The Jackson Laboratory and used for backcrossing of all mouse strains. *Scn2a*^+/-^ mice were provided by Drs. E. Glasscock and M. Montal (Mishra et al., 2017; Planells-Cases et al., 2000). *Scn2a*^*+/fl*^ mice were created as described previously (Spratt et al., 2019). Mice were anesthetized with isoflurane and positioned in a stereotaxic apparatus. 500 nL volumes of AAV-EF1A-Cre-IRES-mCherry (UNC Vector Core) were injected into the mPFC of *Scn2a*^+/+^, *Scn2a*^*fl/+*^ and *Scn2a*^*fl/fl*^ mice (stereotaxic coordinates [mm]: anterior-posterior [AP], +1.7, mediolateral [ML] −0.35; dorsoventral [DV]: −2.6). Mice were used in experiments four weeks post injection.

### *Ex vivo* electrophysiology and two-photon imaging

Two-photon imaging and current- and voltage-clamp electrophysiology were performed as described previously (Spratt et al., 2019). Briefly, mice at postnatal day 56-88 were anesthetized and 250 µm-thick coronal slices containing medial prefrontal cortex were prepared. Slices were prepared from *Scn2a*^+/-^, *Scn2a*^*+/fl*^, Scn2a^fl/fl^, or *Scn2a* wild type littermates (genotyped by PCR). All data were acquired and analyzed blind to *Scn2a* genotype. Data were acquired from both sexes (blind to sex). Cutting solution contained (in mM): 87 NaCl, 25 NaHCO_3_, 25 glucose, 75 sucrose, 2.5 KCl, 1.25 NaH_2_PO_4_, 0.5 CaCl_2_ and 7 MgCl_2_; bubbled with 5%CO_2_/95%O_2_; 4C. Following cutting, slices were incubated in the same solution for 30 min at 33C, then at room temperature until recording. Recording solution contained (in mM): 125 NaCl, 2.5 KCl, 2 CaCl_2_, 1 MgCl_2_, 25 NaHCO_3_, 1.25 NaH_2_PO_4_, 25 glucose; bubbled with 5%CO_2_/95%O_2_; 32-34C, ∼310 mOsm. Neurons were visualized with Dodt-contrast optics for conventional visually guided whole-cell recording, or with 2-photon-guided imaging of reporter-driven tdTomato fluorescence overlaid on an image of the slice (scanning DIC). For current-clamp recordings and voltage-clamp recordings of K^+^ currents, patch electrodes (Schott 8250 glass, 3-4 M tip resistance) were filled with a solution containing (in mM): 113 K-Gluconate, 9 HEPES, 4.5 MgCl_2_, 0.1 EGTA, 14 Tris_2_-phosphocreatine, 4 Na_2_-ATP, 0.3 tris-GTP; ∼290 mOsm, pH: 7.2-7.25. For Ca^2+^ imaging, EGTA was replaced with 250 µM Fluo-5F and 20 µM Alexa 594. For voltage-clamp recordings of persistent Na+ currents, internal solution contained (in mM): 110 CsMeSO_3_, 40 HEPES, 1 KCl, 4 NaCl, 4 Mg-ATP, 10 Na-phosphocreatine, 0.4 Na_2_-GTP, 0.1 EGTA; ∼290 mOsm, pH: 7.22. All data were corrected for measured junction potentials of 12 and 11 mV in K- and Cs-based internals, respectively.

Electrophysiological data were acquired using Multiclamp 700A or 700B amplifiers (Molecular Devices) via custom routines in IgorPro (Wavemetrics) from layer 5b thick tufted neurons, using metrics as previously described (Spratt et al., 2019). Data were acquired at 50 kHz and filtered at 20 kHz. For current-clamp recordings, pipette capacitance was compensated by 50% of the fast capacitance measured under gigaohm seal conditions in voltage-clamp prior to establishing a whole-cell configuration, and the bridge was balanced. For voltage-clamp recordings, pipette capacitance was compensated completely, and series resistance was compensated 50%. Series resistance was <15 M in all recordings. Experiments were omitted if input resistance changed by > ±15%. AP threshold and peak dV/dt measurements were determined from the first AP evoked by a near-rheobase current in pyramidal cells (300 ms duration; 10 pA increments), or the first AP within a train of APs with a minimum inter-AP frequency of 25 Hz in inhibitory neurons. Threshold was defined as the V_m_ when dV/dt measurements first exceeded 15 V/s. AHPs were defined as the minimum voltage between APs.

Persistent Na^+^ and currents were activated with 500 ms voltage steps from −90 mV and corrected using p/n leak subtraction. 10-15 trials were averaged per voltage step. Current amplitudes were calculated as the average of the last 100 ms of each step. Experiments were performed in 25 μM picrotoxin, 10 μM NBQX, 10 mM TEA, 2 mM 4-AP, 200 μM Cd^2+^, 2 μM TTA-P2, and 1 mM Cs^+^. K^+^ currents were activated with 500 ms voltage steps from -90 to -20 mV, in 10 mV increments. 5 trials were averaged per voltage step. Current amplitudes were calculated from the transient peak and sustained components (last 50 ms). Experiments were performed in 500 nM TTX, 25 μM picrotoxin, 10 μM NBQX, and 1 mM Cs^+^. Ca^2+^ channels were not blocked to allow for activation of Ca^2+^-dependent K^+^ channels.

Dynamic clamp experiments were performed using a dPatch amplifier (Sutter Instrument) with identical internal and external solutions as in current-clamp experiments. For these experiments, pipettes were wrapped with parafilm to reduce pipette capacitance to 7.2 ± 0.1 pF, and capacitance was compensated to 90% of values obtained when establishing gigaohm seals. Series resistance (6.6 ± 0.2 MΩ) was bridge balanced completely. Data were acquired at 100-200 kHz and low pass filtered at 10-20 kHz. Na_V_1.2 was simulated with an 8 state Markov model from (Hallermann et al., 2012) and K_V_1.2 was simulated with a “K_t” Hodgkin Huxley-based model from (Hay et al., 2011). In both cases, a voltage offset variable was added to account for junction potential offsets.

### Modeling

A pyramidal cell compartmental model was implemented in the NEURON environment (v7.5), based on a Blue Brain Project model of a thick-tufted layer 5b pyramidal cell (TTPC1) (Markram et al., 2015; Ramaswamy and Markram, 2015), with baseline distributions of Na_V_1.2 and Na_V_1.6 set as previously described (Ben-Shalom et al., 2017). For phase plane comparisons, the first AP evoked with 500 pA stimulus intensity (25 ms duration) were compared in each model configuration, with threshold considered the membrane potential when dV/dt exceeds 15 V/s.

### Immunofluorescence

Tissue samples were collected from P60-90 *Scn2a*^*fl/fl*^ mice stereotaxically injected with pAAV-Ef1a-mcherry-IRES-Cre unilaterally into mPFC after intracardiac perfusion with 4% paraformaldehyde in phosphate buffered saline (PBS). The tissue was then fixed in 4% paraformaldehyde for 2 hours and cryopreserved in increasing concentrations of sucrose in PBS (15% then 30%) overnight at 4°C. Tissue samples were embedded at -80°C in Tissue-Tek O.C.T compound and sectioned coronally into 30-μm thick sections on a cryostat. Sections were rinsed in PBS and then blocked with 10% normal goat serum with 0.2% TritonX-100 in PBS for 40 minutes at room temperature. Sections were incubated overnight with primary antibodies diluted in 0.1% TritonX-100 in PBS with 2% normal goat serum at 4°C. Primary antibodies were as follows: Rabbit IgG Anti-Na_V_1.6 (Alomone ASC-009; 1:200), rabbit IgG Anti-Na_V_1.2 (Abcam ab65163; 1:250), mouse IgG2a Anti-ankyrin-G (Neuromab 75-146; 1:500). Sections were then rinsed in PBS and incubated for 2 hours in secondary antibodies diluted in 0.1% TritonX-100 in PBS with 2% normal goat serum at room temperature. Secondary antibodies were Alexa Fluor 488-conjugated goat anti-rabbit IgG (1:500; A11070, Thermo Fisher Scientific), and Alexa Fluor 647-conjugated goat anti-mouse igG2a (1:500; A21241, Thermo Fisher Scientific). Sections were then rinsed in PBS and slides were mounted with ProLong Gold Antifade mounting media with DAPI (P36931, Thermo Fisher Scientific) and stored at 4°C until imaging.

Images were collected on a Fluoview3000 (Olympus) using appropriate bandpass filters with a 40x 1.4 NA objective with laser intensities and photomultiplier voltages held constant across sections. Serial z-stacks were acquired at 0.1 µm steps in Z, with XY zoom adjusted to 2x Nyquist resolution. Data were stitched using proprietary Olympus software with no adjustment of pixel intensities near borders between stacks. Fluorescent intensity in the AIS was analyzed using image-J by manually drawing a region of interest around the AIS using ankyrin-G labeling as a guide (Na_V_ channel was not visible to the experimentalist). Average pixel intensity was then calculated within this region for Na_V_ and ankyrin-G channels. Analysis was limited to PFC layer 5b using established layer divisions (Clarkson et al., 2017). Na_V_1.6 intensity was normalized to the average ankyrin-G intensity per section to control for variability across animals. Na_V_1.2 staining in the AIS, which was associated with a higher level of diffuse neuropil staining than Na_V_1.6, was first background subtracted from regions of interest of identical size and shape immediately adjacent to each AIS, then normalized to ankyrin-G intensity.

### Chemicals

Fluo-5F pentapotassium salt and Alexa Fluor 594 hydrazide Na^+^ salt were from Invitrogen. Picrotoxin, R-CPP, and NBQX were from Tocris. TTX-citrate was from Alomone. All others were from Sigma.

### Quantification and Statistical Analysis

Data are summarized either with box plots depicting the median, quartiles, and 90% tails or with violin plots with individual datapoints overlaid. n denotes cells for all electrophysiology, spines for spine morphology, and animals for behavior. Data were obtained from 2-14 mice per condition for electrophysiology and imaging experiments. Group sample sizes were chosen based on standards in the field, and no statistical methods were used to predetermine sample size. Unless specifically noted, no assumptions were made about the underlying distributions of the data and two-sided, rank-based nonparametric tests were used. Statistical tests are noted throughout text. Significance was set at an alpha value of 0.05, with a Bonferroni correction for multiple comparisons when appropriate. Statistical analysis was performed using Statview (SAS), and custom routines in MATLAB R2016b (Mathworks), and Python 3.6.4.

---

**Figure S1.**
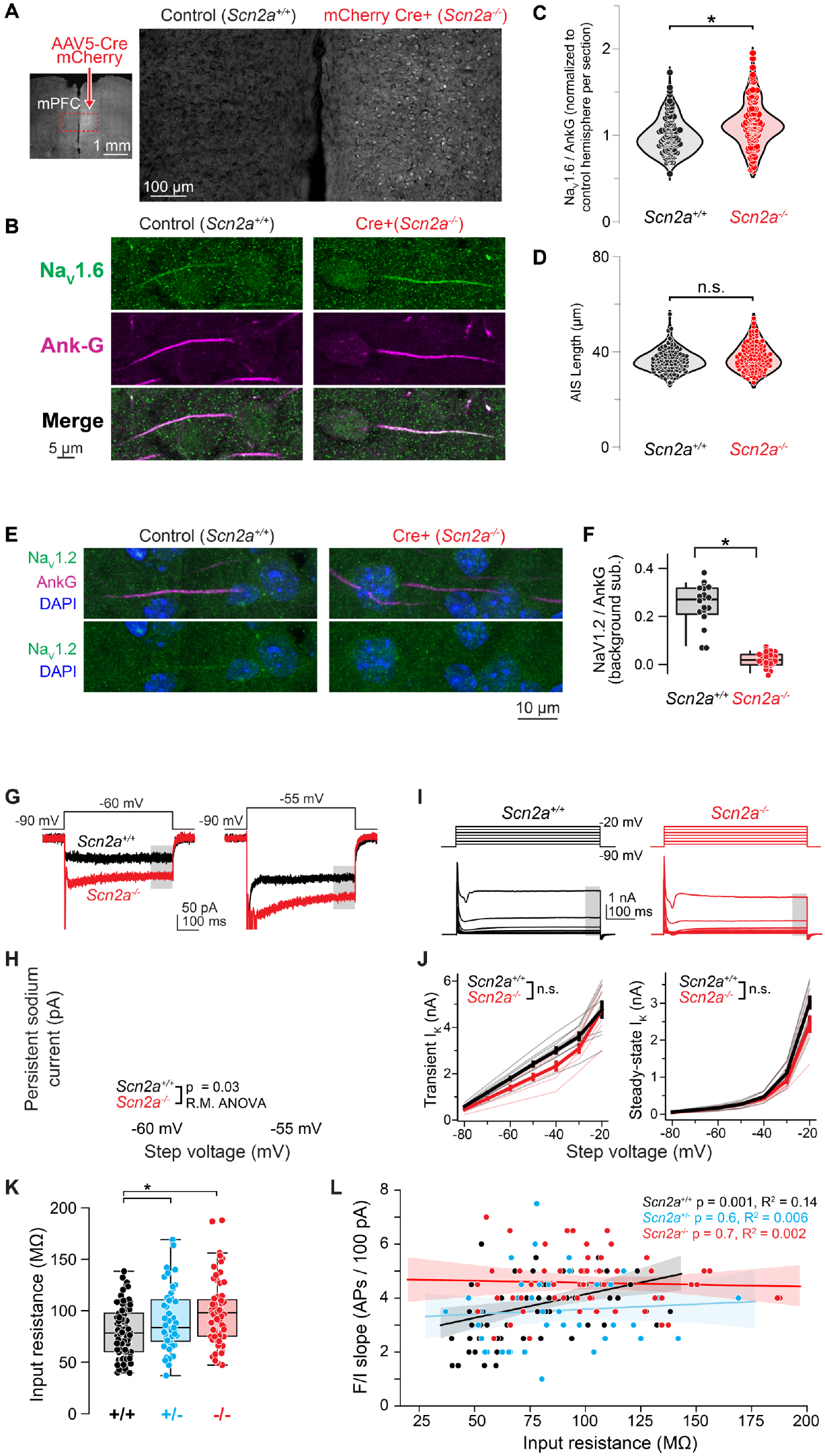
related to Fig. 1: Currents and intrinsic properties of *Scn2a*^*-/-*^ cells. **A:** Example injection of unilateral AAV5-mCherry-Cre into mPFC of mouse at P28. Comparisons were made between injected (*Scn2a*^*-/-*^) and control (*Scn2a*^*+/+*^) hemispheres. **B:** Immunostaining of Na_V_1.6 and ankyrin-G in layer 5 of both hemispheres. **C:** Quantification of Na_V_1.6 staining. Intensity in each AIS was normalized to ankyrin-G. Data were then normalized to mean ankyrin-G intensity of all initial segments analyzed within each section (both hemispheres). Data plotted as violin plots with individual initial segments as single points. *: p < 0.001, Mann-Whitney. n = 126 WT and 123 *Scn2a*^*-/-*^ initial segments across 3 animals. **D:** AIS length was not altered by Na_V_1.2 knockout (p = 0.14, Mann-Whitney). **E:** Immunostaining of Na_V_1.2 and ankyrin-G in both hemispheres. **F:** Quantification of Na_V_ 1.2 staining. n = 20 WT, 24 *Scn2a*^*-/-*^; *: p < 0.0001, Mann-Whitney. Na_V_ 1.2 intensity is not different than 0 (p = 0.07, one sample t-test). **G:** Example persistent sodium currents from *Scn2a*^*+/+*^ (black) and *Scn2a*^*-/-*^ (red) cells for voltage steps from -90 to -60 or -55 mV. **H:** Summary of persistent currents for each voltage step. p = 0.03, repeated measures ANOVA across genotypes. **I:** Example whole cell potassium currents from *Scn2a*^*+/+*^ (black) and *Scn2a*^*-/-*^ (red) cells for voltage steps from -90 to -20 mV in 10 mV increments. **J:** Summary for transient onset current and steady state current (last 50 ms before voltage step offset, grey area in I). Thick lines and bars are mean ± SEM across population. Light, thin lines are data from single cells. Data analyzed with repeated measures ANOVA. **K:** Input resistance for all cells in Fig. 1. *: p < 0.001, Kruskall-Wallis test, Wilcoxon rank sum posthoc test. **L:** Input resistance vs. the slope of the F/I curve between 100 and 300 pA for each cell. Data color-coded as in K. Lines and shaded regions represent best linear fit and 95% confidence intervals. p and R2 values noted in figure.

**Figure S2.**
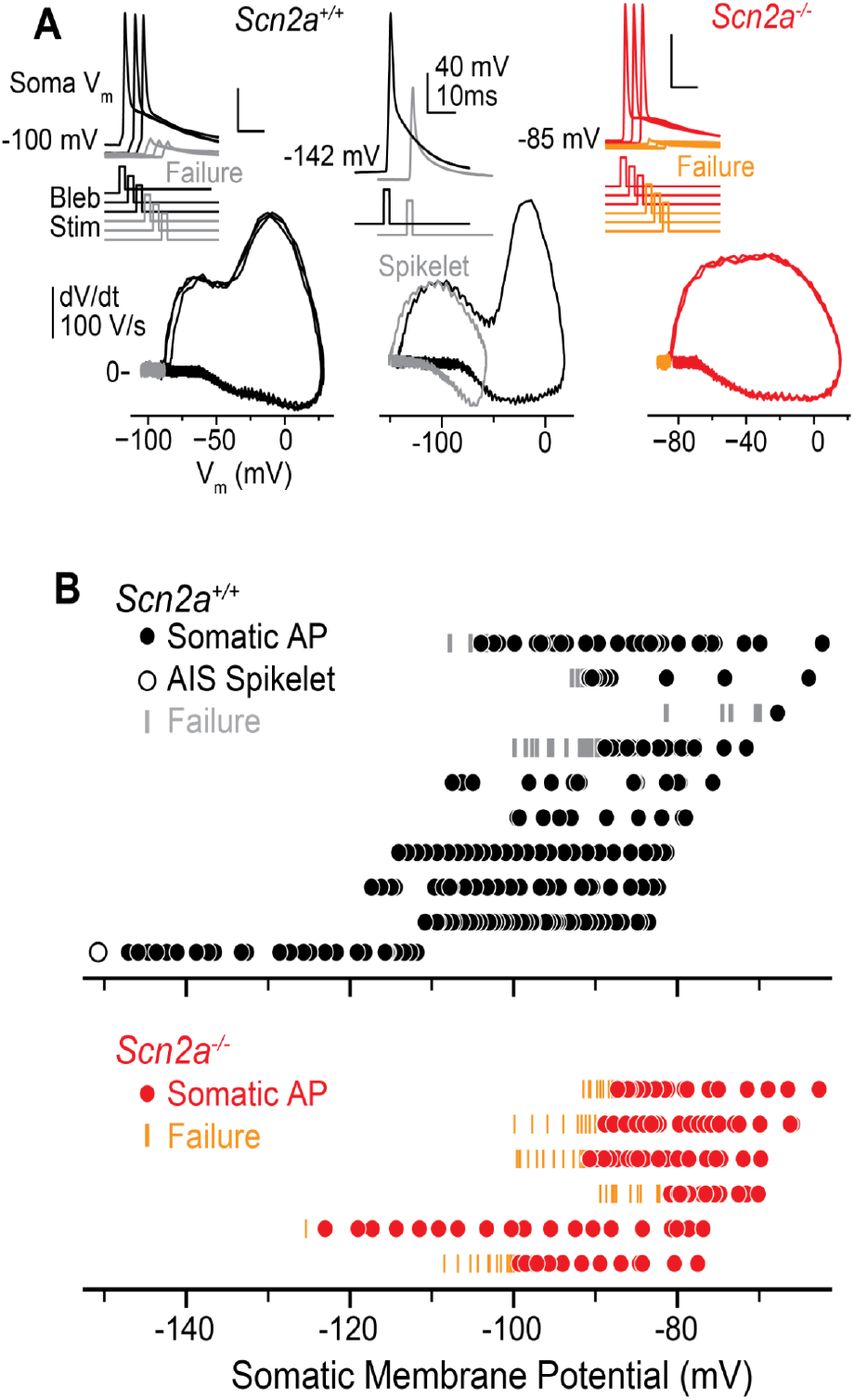
related to Fig. 2: Somatic invasion of APs initiated from axonal bleb. **A:** Examples of successive single APs evoked with 2 ms, 2 nA current injection at bleb. Somatic Vm was hyperpolarized on successive trials with steps of somatic bias current, eventually hyperpolarizing to the point that no APs could be evoked via bleb current injection. Bleb stim timing offset for clarity. Similar effects noted in WT and *Scn2a*^*-/-*^ cells. Spikelet observed once at -150 mV in WT cell (middle). Note differences in X scale for each phase-plane. **B:** Summary across all WT and *Scn2a*^-/-^ cells. Each row is a single cell, with somatic AP successes, failures, and spikelets at each somatic voltage plotted as closed circles, vertical lines, and open circles, respectively.

**Figure S3.**
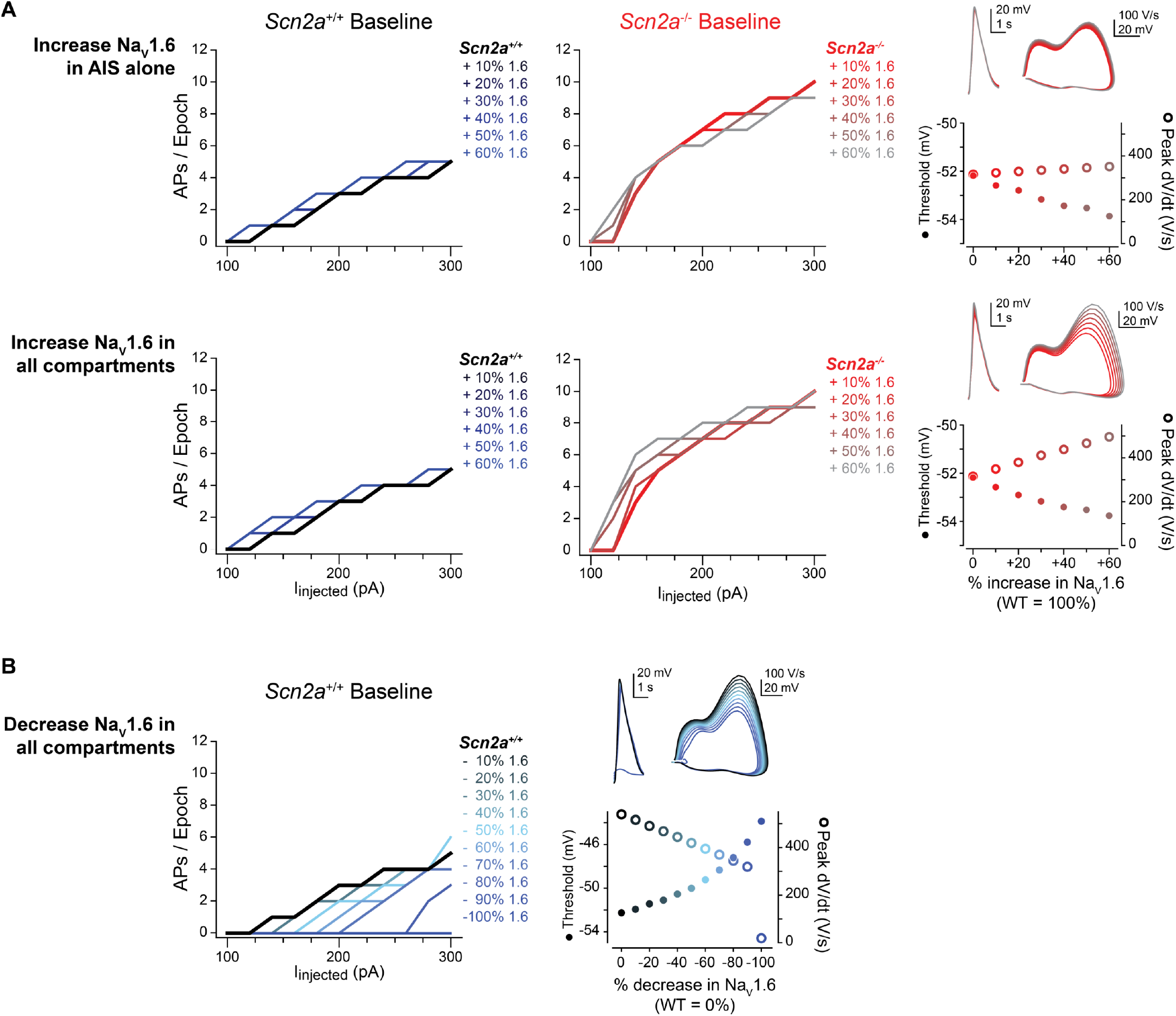
related to Fig. 3: Altering Na_V_1.6 density in models does not recapitulate empirical results. **A:** Increased Na_V_1.6 density (10% increments over baseline levels in each compartment) either in the AIS alone (top row, matching immunofluorescence and persistent current observations) or in all compartments (bottom row, assuming similar effects as those observed in AIS) has minimal effect on F/I curves in WT models (left, blue), or in *Scn2a*^*-/-*^ models (right, red). Note that, in both cases, AP threshold hyperpolarizes with increased Na_V_1.6 density, an effect not observed *ex vivo* (Fig. 1). **B:** Decreased Na_V_1.6 density (10% increments from baseline WT levels to full knockout, all compartments) increases AP threshold and reduces F/I curve. Contrast with Na_V_1.2 knockout (Fig. 3).

**Figure S4.**
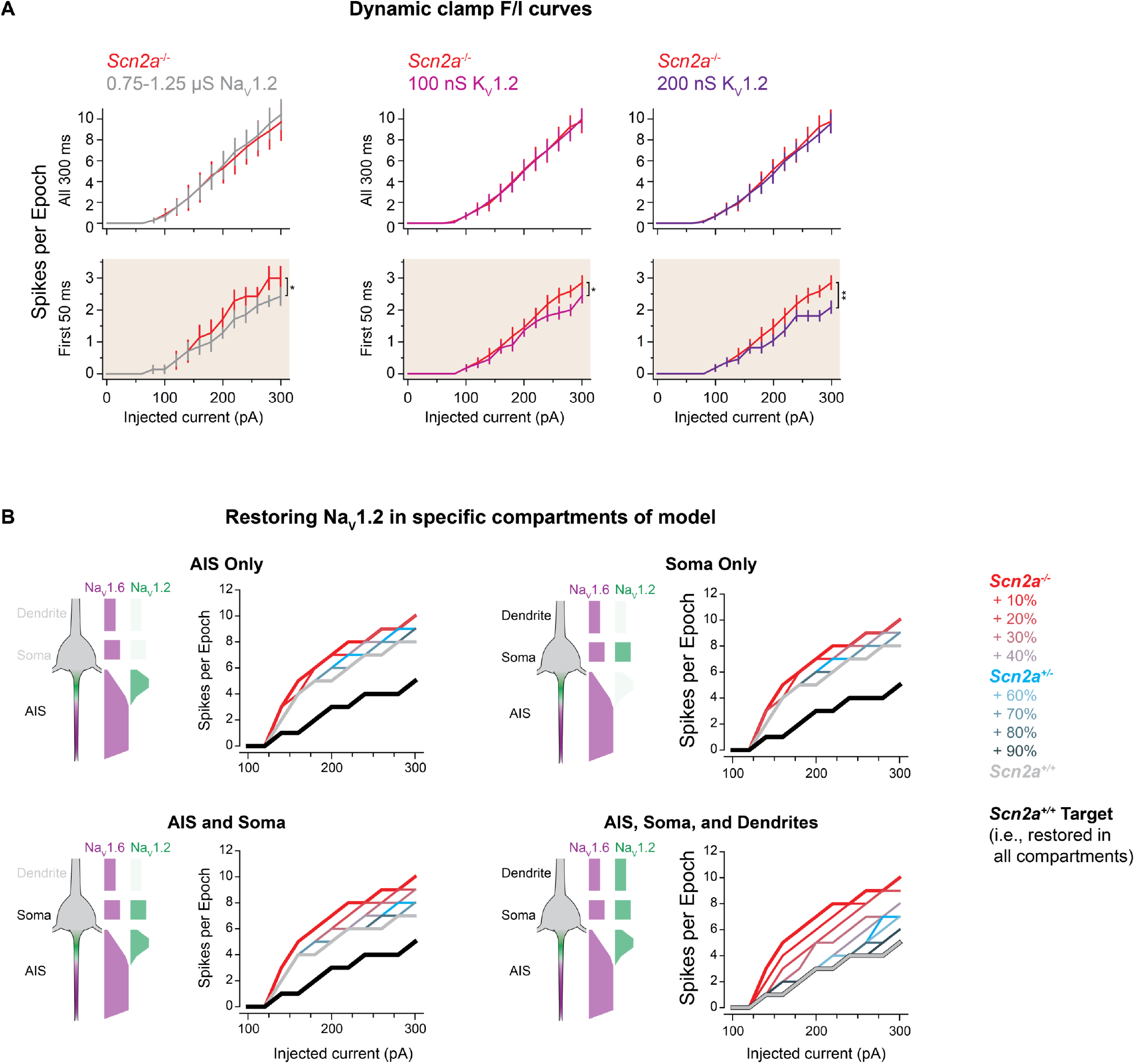
related to Fig. 4: Limitations in space-clamp of injected conductances prevents full restoration of *Scn2a*^*+/+*^-like F/I curves in *Scn2a*^*-/-*^ cells. **A:** F/I curves for data in Fig. 4. Baseline vs. dynamic clamp injection compared in all cases for entire 300 ms epoch or first 50 ms (brown background), as in Fig. 1. Data color-coded as noted above panels. Bars are mean ± SEM. *: p< 0.05, **: p<0.01 for changes in slope between 100 and 300pA, Wilcoxon signed-rank test. **B:** Compartmental models in which Na 1.2 density was progressively restored from *Scn2a*^*-/-*^ to *Scn2a*^*+/+*^ levels (10% increments) in specific compartments. Data compared to “target” of complete restoration in all compartments (black: AIS, soma, dendrite). Note difficulty separating *Scn2a*^*-/-*^ curve from 80% restoration curve in AIS or soma-only conditions.

